# GLMdenoise improves multivariate pattern analysis of fMRI data

**DOI:** 10.1101/320838

**Authors:** Ian Charest, Nikolaus Kriegeskorte, Kendrick N. Kay

## Abstract

GLMdenoise is a denoising technique for task-based fMRI. In GLMdenoise, estimates of spatially correlated noise (which may be physiological, instrumental, motion-related, or neural in origin) are derived from the data and incorporated as nuisance regressors in a general linear model (GLM) analysis. We previously showed that GLMdenoise outperforms a variety of other denoising techniques in terms of cross-validation accuracy of GLM estimates (Kay et al., 2013a). However, the practical impact of denoising for experimental studies remains unclear. Here we examine whether and to what extent GLMdenoise improves sensitivity in the context of multivariate pattern analysis of fMRI data. On a large number of participants (31 participants across 4 experiments; 3 T, gradient-echo, spatial resolution 2–3.75 mm, temporal resolution 1.3–2 s, number of conditions 32–75), we perform representational similarity analysis (Kriegeskorte et al., 2008a) as well as pattern classification (Haxby et al., 2001). We find that GLMdenoise substantially improves replicability of representational dissimilarity matrices (RDMs) across independent splits of each participant’s dataset (average RDM replicability increases from *r* = 0.46 to *r* = 0.61). Additionally, we find that GLMdenoise substantially improves pairwise classification accuracy (average classification accuracy increases from 79% correct to 84% correct). We show that GLMdenoise often improves and never degrades performance for individual participants and that GLMdenoise also improves across-participant consistency. We conclude that GLMdenoise is a useful tool that can be routinely used to maximize the amount of information extracted from fMRI activity patterns.

## 1. INTRODUCTION

Noise is a critical concern when collecting and analyzing functional magnetic resonance imaging (fMRI) data. The blood oxygenation-level dependent (BOLD) signal measured with fMRI contains many sources of noise (e.g., physiological noise, instrumental noise) that are mixed with task-specific signals of interest. In order to draw valid conclusions regarding how a cognitive experiment affects the BOLD signal, it is necessary to extract meaningful signals from the data, limiting various noise contaminations. Classical analyses of fMRI data involve performing a general linear model analysis (Friston et al., 1994, 1995b; Worsley and Friston, 1995). This consists of modelling the time-series of each fMRI voxel using a design matrix that characterizes the onsets of an experiment’s conditions. These events are convolved with a hemodynamic response function (HRF; Boynton et al., 1996; Lindquist et al., 2009), and then a least-squares optimization is performed to minimize the distance between the data and the model. The result of this process is a set of beta weights that characterize fMRI voxel activities.

A common approach to improve the sensitivity of the model is to incorporate nuisance regressors in the design matrix (Friston et al., 1995a; Lund et al., 2006). These nuisance regressors often include participant motion estimates and/or linear and non-linear drift terms, with the goal of improving parameter estimates by accounting for these potential sources of noise (see Ciric et al., 2017 for how nuisance regressors are applied the context of resting-state fMRI). The choice of these regressors is often arbitrary, dependent on the philosophy of the software package used to analyze the data, and may harm task-related estimates if they are inaccurate characterizations of the noise. To address these issues, we previously introduced GLMdenoise (Kay et al., 2013a). GLMdenoise is a denoising technique, inspired by previous work (Behzadi et al., 2007; Bianciardi et al., 2009; Fox et al., 2006), that improves signal-to-noise-ratio (SNR) by automatically deriving the noise regressors entered in the general linear model (GLM) through careful cross-validated analysis of the fMRI time-series. The noise regressors are derived by application of principal components analysis (PCA) on time-series of voxels unrelated to the experimental paradigm, and cross-validation is used to automatically select the appropriate number of regressors for each given dataset. The noise regressors derived in GLMdenoise are general and can encompass many different types of noise, including motion-related noise, physiological noise, and neural noise^1^. A MATLAB toolbox that implements GLMdenoise is freely available at http://cvnlab.net/GLMdenoise/.

In the initial study introducing the technique (Kay et al., 2013a), we showed that GLMdenoise outperforms a variety of other denoising methods on a number of datasets. Our metric of performance was univariate cross-validation accuracy of GLM response-amplitude estimates (beta weights). This criterion quantifies how accurately estimates of beta weights match experimentally observed BOLD time-series data. GLMdenoise computes this metric on a voxel-by-voxel basis and does not make reference to any specific brain regions. Although this approach is rigorous and objective, it remains unclear to what extent GLMdenoise brings practical benefits to experimental studies.

The use of multivariate pattern analysis (MVPA), including pattern classification (Haxby et al., 2001; Haynes and Rees, 2006; Kamitani and Tong, 2005; Kriegeskorte et al., 2006; for a review see Haxby, 2012) and representational similarity analysis (RSA) (Charest et al., 2014; Kriegeskorte et al., 2008b, 2008a; Kriegeskorte and Kievit, 2013; Nili et al., 2014) is growing in popularity in neuroimaging studies. These powerful methods enable the use of fMRI to investigate the information represented in patterns of activity within brain regions. In classical MVPA, a classification algorithm is used to determine whether fMRI response patterns contain information that discriminate different conditions. In RSA, fMRI response patterns are compared between all pairwise experimental conditions to reveal the representational geometry characteristic of a given brain region. These comparisons are assembled in a representational dissimilarity matrix (RDM). These RDMs are useful as they provide some insight into a brain region’s information and reveal the format in which it is represented. This approach is increasingly popular in cognitive neuroscience, as it provides a common ground to relate data from multiple measurement techniques (e.g., electrophysiology, behavior, fMRI, computational models, etc.; see Carlson et al., 2013; Cichy et al., 2016a, 2016b, 2014 for examples and Kriegeskorte and Kievit, 2013 for a review).

In light of this paradigm shift in functional neuroimaging, a high priority for neuroscientists is to optimize data quality before attempting MVPA. A number of studies have combined GLMdenoise with multivariate analyses (for example, see Allen et al., 2018; Charest et al., 2014; Erez et al., 2016), but none have evaluated whether it improves such analyses. Thus, an important question is whether GLMdenoise brings benefits to MVPA and whether the benefits are sufficiently substantial and consistent across studies and participants to justify the complications associated with the integration of GLMdenoise into the analysis. Because GLMdenoise improves the ability of GLM beta weights to generalize to unseen data (Kay et al., 2013a), these beta weights have increased accuracy, and it is reasonable to expect that any subsequent analysis of those beta weights will produce higher quality results. But fMRI data and analyses are complex, and it remains a valuable empirical question whether GLMdenoise in fact improves multivariate analyses and how large the improvement might be (e.g., 1% or 10% increase in percent correct for classification performance).

In this paper, we systematically assess the impact of GLMdenoise on multivariate analyses of a large number of participants compiled from four different experiments. Although these are all visual experiments, we believe the principles underlying the technique will likely generalize to other types of experiments (e.g. auditory, cognitive, motor). The stimuli in the experiments ranged from abstract patterns to images of bodies, faces, places and objects (see Materials and Methods). In addition to their condition-rich designs (the experiments range from 32 conditions to 75 conditions), these experiments all had an ample number of repetitions to perform split-half analyses. Being able to perform split-half analyses is critical for evaluating data reliability and replicability. We analyzed these datasets using pattern classification and representational similarity analyses in regions of interest (ROIs) along the ventral visual stream, which included primary visual cortex, visual word form area, fusiform face area, and human inferior temporal cortex. We observed consistent and substantial improvements to results when using GLMdenoise in the analysis pipeline.

## 2. MATERIALS AND METHODS

### 2.1 Participants and datasets

We collected fMRI data from 31 distinct participants. Informed written consent was obtained from all participants. Experimental protocols were approved by the Stanford University Institutional Review Board, Washington University Human Research Protection Office, and the Cambridge Psychology Research Ethics Committee. Each participant’s dataset consisted of either one or two scan sessions, and each scan session consisted of multiple runs. All fMRI data were collected using a 3T MR scanner and a T2*-weighted, single-shot, gradient-echo pulse sequence with interleaved slice acquisition (see Table 1 for details). Regions-of-interest (ROIs) were defined based on independent localizer data.

**Table 1.** Summary of datasets.

| *Experiment* | *Participants* | *Number of conditions* | *Condition duration (seconds)* | *Number of repetitions per run (or run set)* | *Number of runs (or run sets)* | *Scanner* | *Voxel size (mm)* | *TR (seconds)* | *TE (ms)* | *Flip angle (deg)* | *Volume dimensions* | *Number of volumes per run* | *Total number of runs* | *Data split* |
| --- | --- | --- | --- | --- | --- | --- | --- | --- | --- | --- | --- | --- | --- | --- |
| 1 | 1–3 | 69 | 3 | 1 | 4 | 3T1 | 2.5 | 1.323751 | 29.7 | 71 | 64 x 64 x 21 | 270 | 8 | even/odd |
| 2 | 4–6 | 75 | 3.5 | 1 | 4 | CNI | 2 | 2.006553 | 31 | 77 | 80 x 80 x 26 | 136 | 12 | even/odd |
| 3 | 7–11 | 32 | 4 | 2 | 10 | NIL | 2.5 | 2 | 30 | 77 | 80 x 80 x 28 | 156 | 10 | even/odd |
| 4 | 12–31 | 72 | 1 | 1 | 18 (9 per session) | CBU | 3 x 3 x 3.75 | 2 | 30 | 78 | 64 x 64 x 32 | 216 | 18 | by session |

3T1 = Lucas Center at Stanford University, 3T GE Signa HDX scanner, Nova quadrature or 8-channel RF coil; CNI = Stanford Center for Neurobiological Imaging, 3T GE Signa MR750 scanner, Nova 16-channel RF coil; NIL = Neuroimaging Laboratory at Washington University, 3T Siemens Skyra scanner, Siemens 32-channel RF coil; CBU = Medical Research Council–Cognition and Brain Sciences Unit, 3T Siemens Trio scanner, Siemens 12- channel RF coil. Additional notes: All experiments used a T2\*-weighted, single-shot, gradient-echo pulse sequence, either spiral-trajectory (3T1) or echo-planar imaging (CNI, NIL, CBU).

#### Experiment 1 (3 participants; Participants 1–3)

These data are taken from a previously published study (Kay et al., 2013b) and correspond to datasets 1–3 from the original GLMdenoise paper (Kay et al., 2013a). In this experiment, participants viewed high-contrast black-and-white noise patterns while performing a task at central fixation. Each stimulus condition lasted 3 s, and conditions varied with respect to the visual field location of the noise patterns (we refer to these locations as apertures). There were 69 conditions: 31 vertical apertures (ordered from left to right), 31 horizontal apertures (ordered from bottom to top), and 7 circular apertures (ordered from center to periphery). The ROI for this experiment is primary visual cortex (V1). (Note: this experiment originally consisted of 5 sets of runs; in order to achieve even test-retest splits of the data, we include in this paper only the first 4 sets of runs.)

#### Experiment 2 (3 participants; Participants 4–6)

These data are taken from a previously published study (Kay et al., 2015). In this experiment, participants viewed grayscale faces while performing one of three tasks. Each stimulus condition lasted 3.5 s, and conditions varied with respect to the visual field location of the faces and the task performed. There were 75 conditions: 25 locations (taken from a 5 × 5 grid; ordered from left to right, then top to bottom) × 3 tasks (digit task: one-back task on centrally presented digits, dot task: detection of a dot superimposed on the faces, face task: one-back task on face identity). The ROI for this experiment is fusiform face area (FFA), combining both the posterior fusiform gyrus (pFus-faces/FFA-1) and middle fusiform gyrus (mFus-faces/FFA-2) subdivisions of FFA (Weiner et al., 2014).

#### Experiment 3 (5 participants; Participants 7–11)

In this experiment, participants viewed a variety of grayscale stimuli (e.g., faces, words, texture patterns) while performing a one-back task on the stimuli. Each stimulus condition lasted 4 s (e.g., four 100%-contrast faces presented for 800 ms each with a gap of 200 ms), and there were 32 conditions. The ROI for this experiment is visual word form area (VWFA), defined as word-selective cortex in and around the left occipitotemporal sulcus (Yeatman et al., 2013).

#### Experiment 4 (20 participants; Participants 12–31)

These data are taken from a previously published study (Charest et al., 2014). In this experiment, participants were paired and each participant viewed images that were familiar to each participant in a pair (18 images per participant, consisting of bodies, faces, places, and man-made objects) as well as 36 object images (common to all participants). There were 72 conditions in total. Each stimulus condition lasted 1 s, and participants performed an anomaly-detection task indicating whether the stimulus had been subtly changed. The ROI for this experiment is human inferotemporal cortex (hIT), which is defined as a wide expanse of posterior and anterior temporal cortex, including fusiform face area, lateral occipital complex, and parahippocampal place area.

In each experiment, conditions were presented in random or pseudorandom order within each run, and rest periods were included between conditions and at the beginning and end of each run. In some participants (Participants 7–31), every condition was presented at least once during each run. In other participants (Participants 1–6), conditions were split across multiple runs. For example, Participants 4–6 involved 75 conditions which were split across three runs, each containing 25 conditions; together, the three runs comprise a run set and multiple run sets were collected over the course of the scan session. The specific characteristics of each participant’s dataset are detailed in Table 1.

### 2.2 Data pre-processing

The first five volumes (Experiments 1–2) or eight volumes (Experiment 4) of each run were discarded to allow longitudinal magnetization to reach steady-state. Differences in slice acquisition times were corrected using sinc interpolation (Experiments 1–4). Measurements of the static magnetic field (*B*_0_) were used to correct volumes for spatial distortion (Experiments 1–3). Motion correction was performed using SPM (Experiments 1–4). Final data interpolation occurred at the original voxel resolution (Experiments 1–2, 4) or the resolution of the cortical surface reconstruction generated by FreeSurfer based on T1-weighted anatomical data (Experiment 3).

### 2.3 Summary of GLMdenoise procedure

A summary of the major steps in GLMdenoise is provided here (for full details, please see Kay et al., 2013a). We start with a baseline GLM that includes task regressors capturing effects related to the experiment and polynomial regressors capturing low-frequency drift. A procedure to estimate the HRF is performed, and the accuracy of the GLM is quantified using leave-one-run-out cross-validation. Voxels whose cross-validated *R*^2^ values are less than 0% are then considered for the noise pool. Note that this selection is not tailored to any specific contrast or effect that might exist in the data, but simply assesses whether any of the task regressors produce non-zero variance in the time series for a given voxel. Moreover, even if the noise pool contains voxels of interest, it is still possible to improve GLM estimates for such voxels (Kay et al., 2013a). The noise pool is further refined to brain voxels by discarding voxels whose mean signal intensity fall below one half of the 99th percentile of mean signal intensity values across all voxels. Next, we extract the time-series data observed for voxels in the noise pool, project out the polynomial regressors, and perform principal components analysis (PCA) on these time series. We add the PCs in decreasing order of variance explained to the GLM as nuisance regressors, and systematically evaluate cross-validation performance as a function of the number of PCs added. Finally, the optimal number of PCs is selected (based on median cross-validated *R*^2^ performance across task-related voxels) and used to obtain the final response-amplitude estimates (beta weights).

### 2.4 General linear model (GLM) analysis

For each participant, runs were split into two groups using either an even/odd split (Experiments 1–3) or a by-session split (Experiment 4). Each group of runs was analyzed using GLMdenoise (http://cvnlab.net/GLMdenoise/), with denoising enabled (optimal number of PCs added to the GLM) and denoising disabled (no PCs added to the GLM). A few notes on the application of GLMdenoise to our datasets: Although this paper describes results for specific regions of interest, the denoising itself was not tailored in any way to these regions but was applied to each dataset in its entirety (as is the default). Regarding the choice of HRF, we used the default ‘optimize’ option, indicating that the HRF is estimated from the data. Since the denoising procedure occurs after HRF estimation (see Section 2.3), the denoised and undenoised results reflect a common HRF. Finally, the entire GLMdenoise procedure (including the internal use of cross-validation) was applied independently to each split of the data; this strict splitting ensures that no improper “parameter sharing” or “data peeking” occurred.

To prepare the data for multivariate pattern analysis, beta weights from the GLM analysis were converted to *t* units (Misaki et al., 2010) by dividing each voxel’s beta weights by the square root of the average squared standard error. The final result of the GLM analysis is a set of four matrices containing multivoxel activity patterns. These matrices are denoted **P_*i,d*_** where *i* = 1 (first data split, referred to as *Test*) or 2 (second data split, referred to as *Re-test*) and *d* = 1 (denoising disabled, referred to as *Baseline*) or 2 (denoising enabled, referred to as *Denoised*). Each **P_*i,d*_** has dimensions *v* × *c* where *v* is the number of voxels in the region of interest and *c* is the number of experimental conditions.

To test GLMdenoise against the common practice of including motion parameters in the GLM design matrix, we also fit two additional GLMs. The first GLM extends the baseline GLM by including the 6 rigid-body transformation parameters obtained from motion correction (*x, y, z, pitch, roll, yaw*). The second GLM includes not only the rigid-body parameters but also their squares, their temporal derivatives, and the squares of the temporal derivatives (Friston et al., 1996), yielding a total of 24 additional parameters in the GLM design matrix. The beta weights produced by these two additional GLMs are compared to those produced by GLMdenoise using the multivariate analyses detailed below.

### 2.5 Representational similarity analysis (RSA)

For conceptual background and further details on RSA, we refer the reader to other papers (Charest et al., 2014; Kriegeskorte et al., 2008a; Kriegeskorte and Kievit, 2013; Nili et al., 2014). Each set of multivoxel activity patterns **P_*i,d*_** was converted into a representational dissimilarity matrix (RDM) **R_*i,d*_** using one minus Pearson’s correlation (*r*) as the metric of dissimilarity. Specifically, the element in the *m*th row and *n*th column of **R_*i,d*_** was computed as one minus the correlation between the *m*th and *n*th columns of **P_*i,d*_**. Each **R_*i,d*_** has dimensions *c* × *c* where *c* is the number of experimental conditions. We then computed three metrics of RDM replicability. *Baseline* is Pearson’s correlation between the lower triangles of **R_1,1_** and **R_2,1_**, and indicates RDM replicability when there is no denoising of either split of the data. *Denoised* is Pearson’s correlation between the lower triangles of **R_1,2_** and **R_2,2_**, and indicates RDM replicability when there is denoising of both splits of the data. *Baseline/Denoised* is the average of Pearson’s correlation between the lower triangles of **R_1,1_** and **R_2,2_** and Pearson’s correlation between the lower triangles of **R_1,2_** and **R_2,1_**, and indicates how well a denoised RDM from one split of the data correlates with an undenoised RDM from the other split of the data. Results are very similar when using Spearman’s correlation as the measure of RDM replicability (data not shown).

To calculate error bars and statistical significance, a bootstrapping procedure was used. In this procedure, a bootstrap sample is constructed by resampling experimental conditions with replacement. The resulting activity patterns are used to compute RDMs, separately for each split of the data. These bootstrapped RDMs are then compared using Pearson’s correlation (as detailed above). The process is repeated 1,500 times to assess variability due to sampling of experimental conditions. *P*-values are calculated by quantifying the fraction of bootstrap samples where a given comparison of interest does not hold (for example, the fraction of cases where Denoised does not lead to higher replicability than Baseline). Note that when calculating RDM metrics, artifactual zeros in RDMs are ignored (multiple instances of the same condition lead to zero distances).

### 2.6 Cross-validated Mahalanobis distance

Recently, multivariate noise-normalized cross-validated Mahalanobis distance (crossnobis) has been proposed as a novel method for computing RDMs (Walther et al., NeuroImage 2016; Diedrichsen et al. PloS Biol 2017). Crossnobis provides unbiased estimates of distances, and can be viewed as an alternative approach to denoising in the context of RSA. In brief, noise covariance between voxels is estimated from GLM residuals using a shrinkage-based estimator (Ledoit and Wolf, 2004) and used to whiten regression coefficients (beta weights), and distances are estimated on an independent partition of the data.

To evaluate the crossnobis method, we performed two analyses. The first analysis involved computing cross-validated Mahalanobis distances based on the design matrix of the baseline GLM (no PCs). This enables assessment of the replicability of RDMs when computed using crossnobis distances. The second analysis involved computing cross-validated Mahalanobis distances based on the design matrix of the final GLM identified by GLMdenoise (optimal number of PCs). This provides an assessment of whether the crossnobis method can benefit from the nuisance regressors identified by GLMdenoise. Note that in these analyses, strict data splitting is observed: the cross-validation used in the crossnobis procedure is fully confined within each data split.

### 2.7 Classification analysis

We performed a simple correlation-based classification analysis, similar to that in (Haxby et al., 2001). Let *m* and *n* refer to indices of two distinct experimental conditions. For both *d* = 1 (Baseline) and *d* = 2 (Denoised), we computed the 2 × 2 matrix **C_*d*_** consisting of Pearson’s correlations between activity patterns, where the rows correspond to activity patterns for conditions *m* and *n* from the first split of the data (the *m*th and *n*th columns of **P_1,*d*_**) and the columns correspond to activity patterns for conditions *m* and *n* from the second split of the data (the *m*th and *n*th columns of **P_2,*d*_**). If the diagonal elements of **C_*d*_** are larger than the off-diagonal elements, this indicates that conditions *m* and *n* can be well discriminated. To convert **C_*d*_** to a single number representing percent correct, we assess whether element (1,1) is greater than (2,1) and whether (2,2) is greater than (1,2) (this treats the first split as the training data and the second split as the testing data) as well as whether element (1,1) is greater than (1,2) and whether (2,2) is greater than (2,1) (this treats the second split as the training data and the first split as the testing data). Percent correct is calculated as the proportion of these four cases where a successful outcome is observed (i.e. diagonal element larger than off-diagonal element). We performed this procedure for every pair of experimental conditions, and then averaged performance across pairs. This yields a single number (pairwise decoding accuracy) that indicates how well conditions can be discriminated from one another.

To assess the statistical significance of the difference in accuracy between Baseline and Denoised, we performed a permutation test in which undenoised and denoised activity patterns are randomly swapped. Specifically, each column of **P_1,1_** is swapped with the corresponding column of **P_1,2_** with probability 0.5, and this procedure is repeated for the columns of **P_2,1_** and **P_2,2_**. After random swapping, differences in accuracy are computed just as in the original procedure. The *p*-value is taken to be the fraction of permutation iterations that exhibit a difference in accuracy that is equal to or larger than the observed difference.

### 2.8 Simulations

We conducted a set of simulations (code available at http://osf.io/bf736) in order to clarify the principles underlying the approach of modeling task-correlated noise (see Supplementary Figure 2). These simulations involved a simple regression model: **y = Xh + Nk + e** where **y** is time-series data (*t* × 1), **X** is a task regressor (*t* × 1), **h** is a task weight (1 × 1), **N** is a nuisance regressor (*t* × 1), **k** is a nuisance weight (1 × 1), and **e** is a set of residuals (*t* × 1). Each simulation consisted of the following steps: (1) Generate task and nuisance regressors by randomly sampling numbers from the normal distribution (*t* = 50), low-pass filtering the resulting values (at a cutoff of 5 cycles per time-series), and *z*-scoring each regressor. The rationale for low-pass filtering is to mimic the properties of fMRI data and to make it more likely that large correlation values between task and nuisance regressors are observed. Two sets of task and nuisance regressors are generated: one set is for training data and the other set is for testing data. (2) Compute the correlation (Pearson’s *r*) between the task and nuisance regressors in the training set. This correlation value is used to bin simulation results. (3) Generate time-series data. We set the true task weight **h** to 100. We control the strength of the nuisance effects by appropriate setting of **k** (e.g., a ‘nuisance level’ of 10 means to use **k** = 1,000 which is 10 times the size of the task weight). We generate residuals **e** by sampling numbers from the normal distribution and scaling the resulting values appropriately (e.g., a ‘residual level’ of 10 means to scale the values by 1,000, which is 10 times the size of the task weight). (4) Fit the time-series data in the training set using two different models. One model consists of just the task regressor **X**. The other model consists of both the task regressor **X** and the nuisance regressor **N**. In both cases, we obtain an estimate of the task weight **h**. (5) For each model, we quantify the accuracy of the estimated **h** by computing the absolute deviation from the true task weight. Results are shown in Supplementary Figure 2A. We also compute cross-validated *R*^2^ as the percentage of variance in the testing data that is explained by the task estimate (task regressor **X** scaled by the estimated **h**). Results are shown in Supplementary Figure 2B.

## 3. RESULTS

We examined to what extent GLMdenoise improves sensitivity in the context of multivariate pattern analysis of fMRI data. To this end, we collected and analysed datasets from 31 fMRI participants. Each dataset involved a large number of experimental conditions presented multiple times over the course of the experiment. We split each dataset into halves (test and re-test) and applied a GLM analysis to each half with and without the use of GLMdenoise. The beta weights returned by the GLM analyses were then assessed using representational similarity analysis (RSA) and pattern classification.

### 3.1 Representational similarity analysis: GLMdenoise improves the replicability of representational dissimilarity matrices

We obtained representational dissimilarity matrices by correlating the pattern of beta weights obtained for each experimental condition (Figure 1). For individual participants, the denoising appears to lead to clearer representational structure shared between test and re-test results (see Supplementary Figure 1 for results on all participants). To further visualize GLMdenoise’s impact on the similarity structure of the RDMs, we applied classical multi-dimensional scaling (MDS; metric stress) to the RDMs obtained from Participant 18 (Figure 2). MDS for the test and re-test RDMs after denoising (bottom row) shows clearer categorical structure (the categories are depicted with red, orange, blue, and cyan circles) than MDS of the test and re-test RDMs when no denoising was performed (top row). In addition, the right column shows that denoising reduces the displacement (or error) of each stimulus in the representational space.

**Figure 1.**
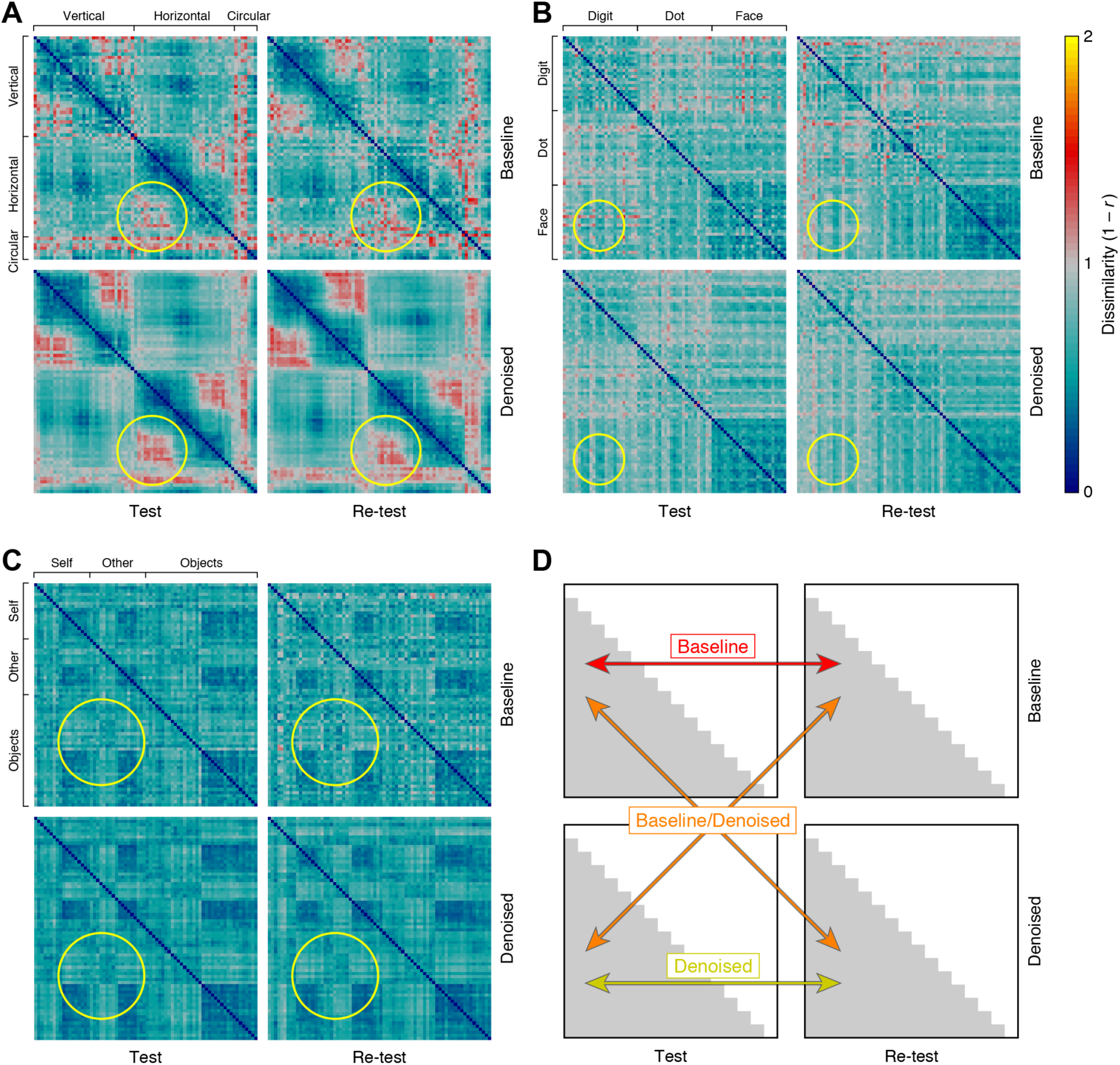
GLMdenoise improves the quality of representational dissimilarity matrices (RDMs). Each participant’s fMRI dataset was split into two halves (Test, Re-test), and each half was analyzed with a standard GLM (Baseline) or with GLMdenoise (Denoised). The results of each analysis are used to construct an RDM, which indicates the dissimilarity between the multivoxel activity patterns associated with each pair of experimental conditions. (A) Results for a participant from Experiment 1 (Participant 3). The yellow circle highlights a section of the RDMs for which denoising yields clearer structure and improves replicability across splits of the data. (B) Results for a participant from Experiment 2 (Participant 4). Same format as A. (C) Results for a participant from Experiment 4 (Participant 26). Same format as A. (D) Metrics of RDM quality. We computed three metrics quantifying the replicability of the lower triangles of the RDMs. *Baseline* is the test-retest correlation of the undenoised RDMs. *Denoised* is the test-retest correlation of the denoised RDMs. *Baseline/Denoised* is the correlation between an undenoised RDM and a denoised RDM (correlation values for the two possible cases are averaged). See Supplementary Figure 1 for RDM results for all participants.

**Figure 2.**
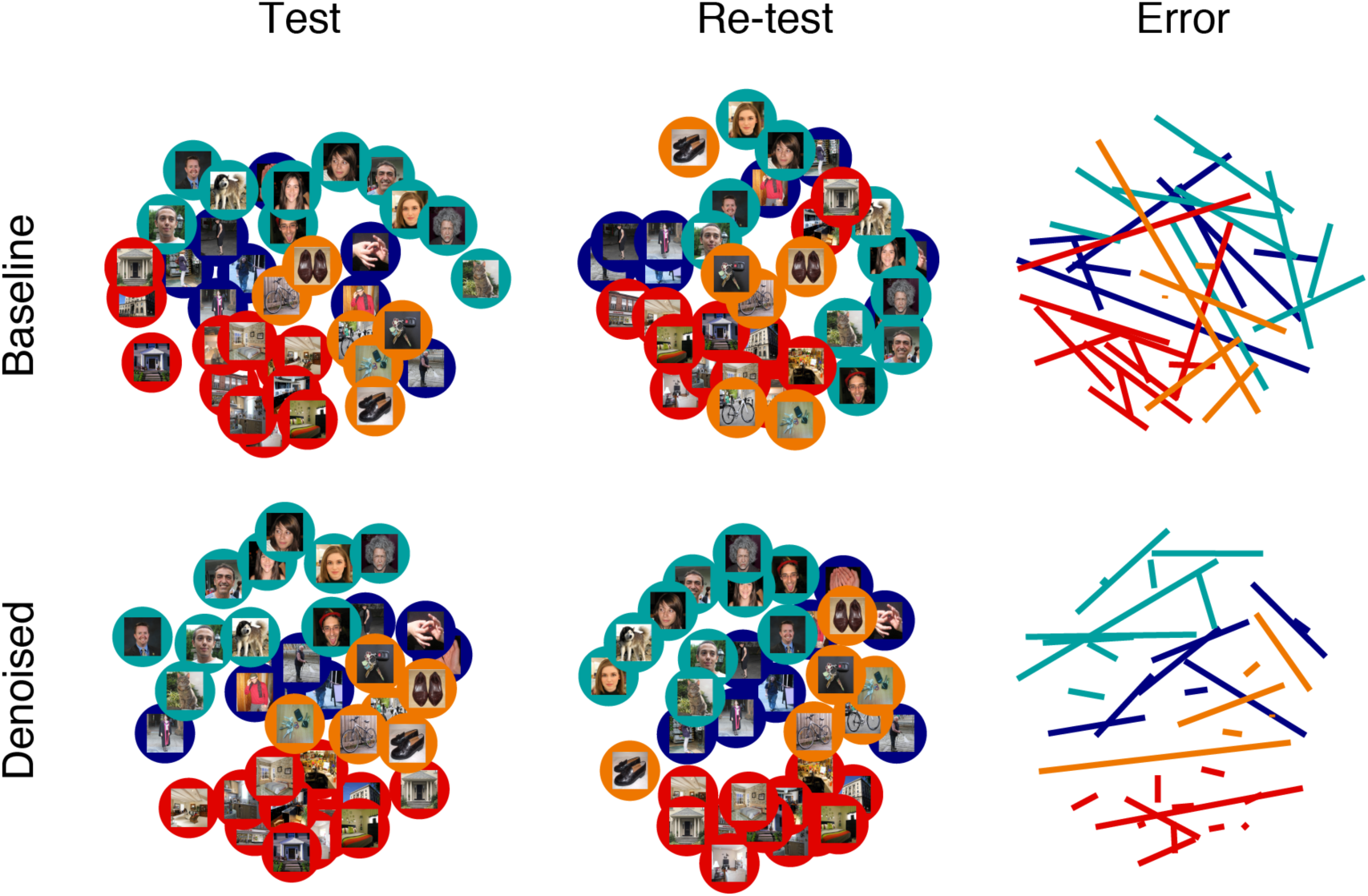
Multidimensional scaling (MDS) illustrates the benefit of denoising RDMs. We apply classical MDS to visualize the similarity structure of the RDMs obtained for Participant 18. Each point is color-coded with regards to stimulus category (red: places, orange: objects, blue: bodies, cyan: faces) and shows the actual stimulus presented to the participant. To quantify the replicability of MDS results across Test and Re-test, we co-registered the two sets of results by first normalizing the scale of each MDS result and then rotating the Re-test MDS result to minimize the error with respect to the Test MDS result. The colored lines depict the distance between Test and Re-test results. Short lines indicate that an item’s position in the representational space is stable across data halves. Long lines suggest that the underlying activity patterns contain noise that is corrupting the RDM results. Denoising substantially improves test-retest replicability of MDS results and increases clustering of similar points. Note that this is just an illustrative example suggesting the potential impact of denoising on neuroscientific results; systematic evaluation of denoising performance for all participants and experiments is provided in later figures.

To further quantify these effects, we computed three performance metrics for all participants. We compared the replicability of the test and re-test RDMs constructed without denoising (Baseline), denoising both data-splits (Denoised), and denoising only one data-split (Baseline/Denoised). Plotting these metrics, we see that RDM replicability is substantially higher when using GLMdenoise (Denoised) compared to baseline (Baseline). In several cases, there are very sizable improvements (Figure 3A), increasing from correlations near 0 to modest correlation values. The high variability in the amount of improvement obtained by denoising is consistent with earlier observations (Kay et al., 2013a) and reflects the fact that the amount of overlap between noise effects and experimental effects depends on a variety of factors such as the number of conditions, their temporal ordering, and incidental differences across participants (such as the amount of head motion).

**Figure 3.**
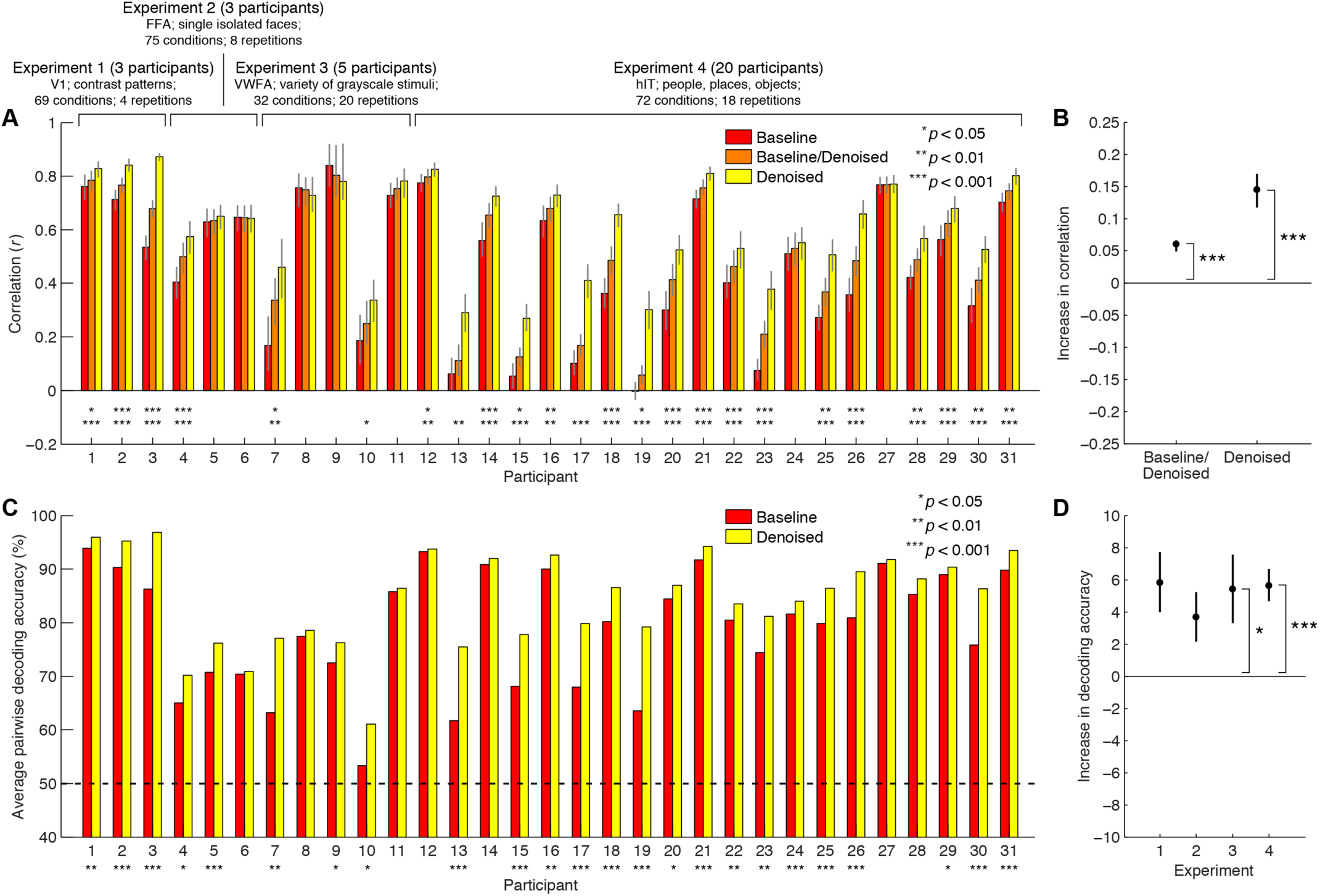
GLMdenoise consistently improves RDM quality and classification performance. (A) RDM metrics for each participant. Error bars (68% confidence intervals) and statistical significance levels were calculated by bootstrapping experimental conditions. The top row of asterisks indicates *p*-values for Baseline/Denoised > Baseline, while the bottom row of asterisks indicates *p*-values for Denoised > Baseline. Many participants exhibit a statistically significant increase in performance relative to Baseline. Importantly, no participant exhibits a statistically significant decrease in performance relative to Baseline. (B) Summary of RDM metrics. For each participant, we compute the increase in correlation relative to Baseline. The median increase across participants is shown (error bars and statistical significance were computed by bootstrapping participants). (C) Classification performance for each participant. The asterisks indicate *p*-values for Denoised > Baseline (permutation test; see Methods). Many participants exhibit a statistically significant increase in performance, and no participant exhibits a decrease in performance. (D) Summary of classification performance. For each participant, we compute the increase in performance relative to Baseline. The mean increase across participants (separated by experiment) is shown. Error bars (standard errors) and statistical significance (*t*-test) were computed parametrically due to the small number of participants in Experiments 1–3.

It is possible that improved replicability reflects biased RDMs. For example, one can imagine a procedure that artificially creates RDM values that are all biased towards 0. Such a procedure would yield enhanced replicability values, but would yield inaccurate RDMs. The Baseline/Denoised metric indicates how well a denoised RDM matches an undenoised RDM from a separate split of the data. Across participants, this metric is higher than Baseline (Figure 3B). This establishes an important control: a raw (undenoised) RDM is better predicted by a denoised RDM compared to another raw RDM. This provides evidence that denoising does not induce bias to RDM structure, but rather, denoises the RDM structure, pushing it closer to the true RDM structure.

### 3.2 Representational similarity analysis: Comparing GLMdenoise to other popular methods

We compared improvements in RDM replicability provided by GLMdenoise to that provided by other denoising methods. One popular method is to include motion parameters in the GLM design matrix; we evaluated a version of this method that involves 6 regressors (rigid-body motion parameters) and a version that involves 24 regressors (rigid-body parameters plus squares and temporal derivatives). We observe an increase in RDM replicability compared to Baseline when including motion parameters (Figure 4A, second and third columns), but the improvements are not as large as those observed under GLMdenoise (Figure 4A, first column).

**Figure 4.**
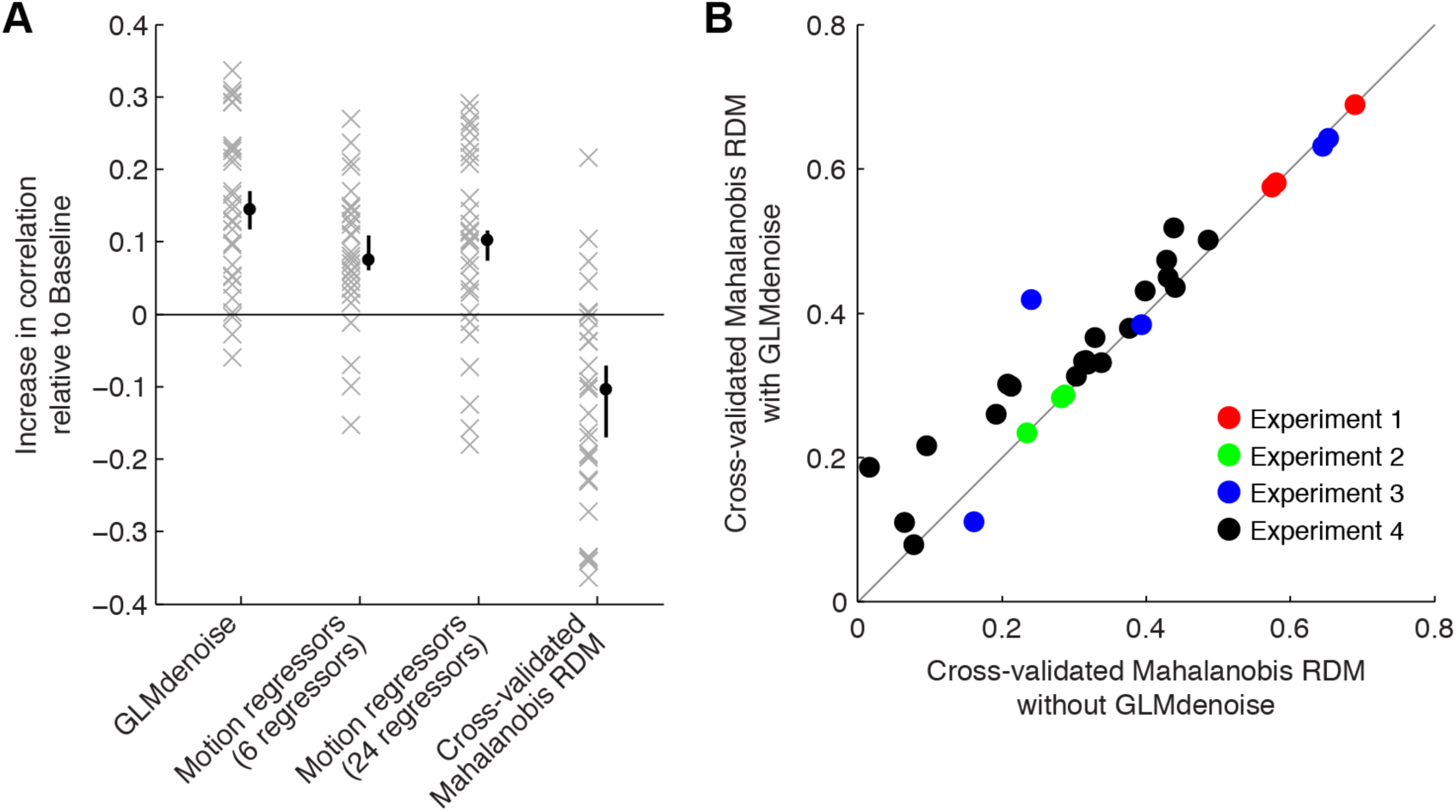
Comparison of GLMdenoise to other approaches. We evaluated the potential benefits of using GLMs augmented with motion parameter estimates and of constructing RDMs using cross-validated Mahalanobis distances (Diedrichsen and Kriegeskorte, 2017; Walther et al., 2016). (A) Summary of results. Each ‘x’ indicates for one participant the increase in RDM replicability (correlation between split-half RDMs) observed relative to Baseline. The black dot indicates the median increase across participants (error bars indicate 68% confidence intervals obtained by bootstrapping participants). In terms of statistical significance (two-tailed bootstrap test), we find that GLMdenoise yields significant improvements compared to Motion regressors (6 regressors) and Cross-validated Mahalanobis RDM (*p* < 0.01) and marginally significant improvements compared to Motion regressors (24 regressors) (*p* = 0.08). For Cross-validated Mahalanobis RDM, the decrease in performance relative to Baseline is statistically significant (*p* < 0.01). (B) GLMdenoise can be combined with cross-validated Mahalanobis distances. We calculated the replicability of RDMs constructed with the cross-validated Mahalanobis distance method, either using the GLM design matrix in the Baseline model (*x*-axis) or using the GLM design matrix provided by GLMdenoise (*y*-axis). Each dot corresponds to one participant. The results show that GLMdenoise is compatible with cross-validated Mahalanobis distance and yields improvements in RDM replicability (two-tailed sign test, *p* < 0.01).

We also assessed RDM replicability for RDMs constructed using cross-validated Mahalanobis distance (Diedrichsen and Kriegeskorte, 2017; Walther et al., 2016). This approach provides unbiased distance estimates and accounts for fMRI noise structure (see Methods). RDM replicability using cross-validated Mahalanobis distance does not show improvements compared to Baseline (Figure 4A, fourth column). The lack of improvement might be because estimation of noise covariance and calculation of distance on independent data require large amounts of data to achieve robust results. Alternatively, Euclidean distance might be a less stable metric than correlation-based distance. We explored an analysis in which the noise regressors identified by GLMdenoise are incorporated into the procedure for computing cross-validated Mahalanobis distance. We find that this combination strategy works well: introducing GLMdenoise yields improvements to RDM replicability (Figure 4B).

### 3.3 Representational similarity analysis: GLMdenoise improves consistency across participants

Thus far, we have only shown that GLMdenoise improves RDM replicability within participants. Do these benefits extend to improved replicability across participants? For each participant, we calculated a single RDM that reflects all data collected for that participant and then calculated Pearson’s correlation between all pairs of participants within each experiment. We find that the majority of pairwise participant comparisons are improved when using GLMdenoise (Figure 5A) and that these improvements are greater than those observed using the motion-parameter approach (Figure 5B). This indicates that not only does GLMdenoise improve the quality of results for individual participants, but these improvements also translate to reduced variability at the group level, thereby enhancing the ability to make inferences about generalization of representational geometries across participants.

**Figure 5.**
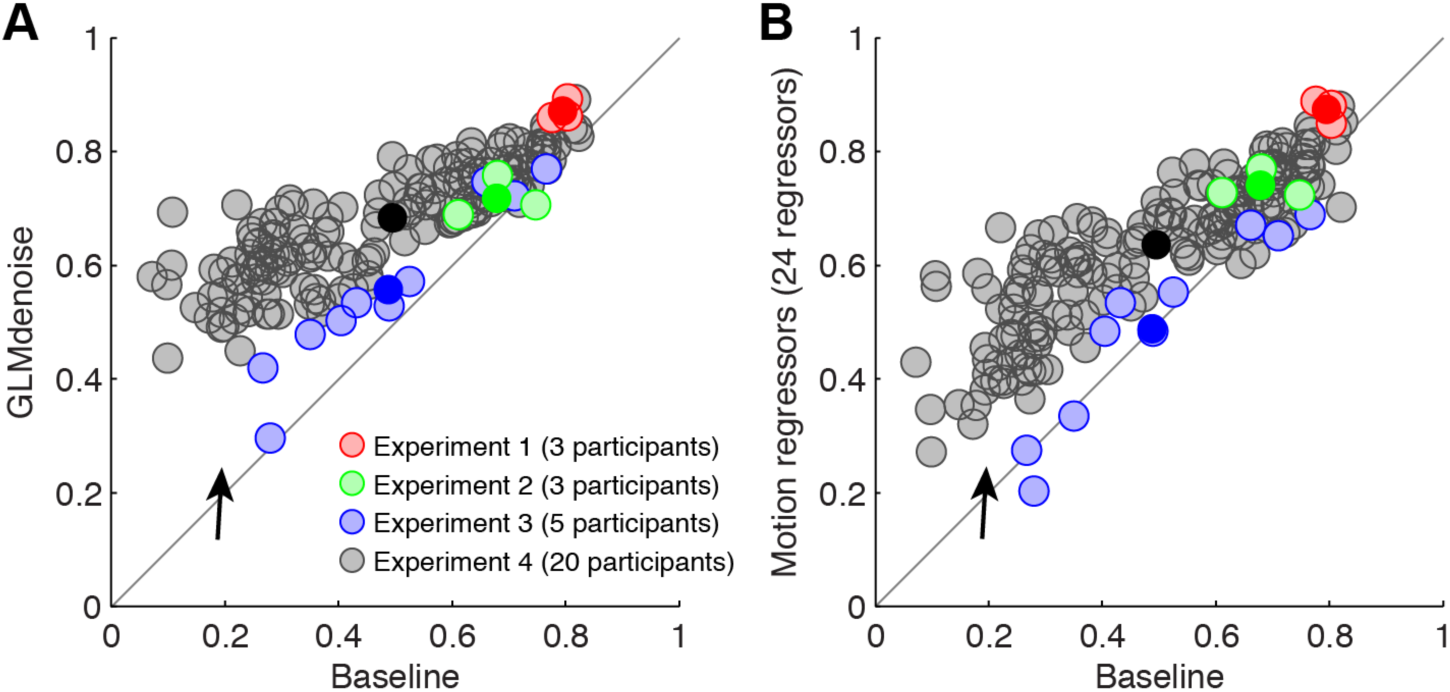
GLMdenoise improves across-participant consistency of representational geometries. For each participant, we averaged the split-half activity patterns to obtain a single set of activity patterns, computed RDMs from these activity patterns, and then computed RDM-to-RDM correlations between all pairs of participants. (For Experiment 4, we restricted this analysis to the 36 of the 72 conditions that were common across participants.) (A) Improvements observed using GLMdenoise. Results with no denoising (*x*-axis) are compared against results with GLMdenoise (*y*-axis). Each light dot indicates one pairwise correlation, and each dark dot indicates the centroid of all pairwise correlations observed for a given experiment. (B) Improvement observed using Motion regressors (24 regressors). Same format as panel A. Overall, GLMdenoise and, to a lesser degree, motion regressors improve across-participant consistency. The black arrow highlights cases in which improvements are especially large. This is consistent with earlier observations that denoising appears to be especially helpful for datasets where signal quality is initially low (see Figure 3).

### 3.4 Pattern classification analysis: GLMdenoise improves decoding accuracy

We have established, using RSA, that multivariate similarity structure has better replicability when using GLMdenoise. If GLMdenoise improves replicability of representational structure, does this also translate into better decoding accuracies between experimental conditions? We performed a complementary analysis that quantifies how well conditions can be discriminated from one another based on fMRI activity patterns, in line with classical MVPA approaches.

We compared the average pairwise decoding accuracy before and after the use of GLMdenoise (see Methods for details). All participants exhibit an increase in decoding accuracy, and in 24 of 31 participants, this increase is statistically significant at *p* < 0.05 (Figure 3C). Since discriminability may be dependent on the specific conditions used in each of our four experiments, we computed the average increase in decoding accuracy independently for each experiment (Figure 3D). We find that in all four experiments, GLMdenoise provides substantial improvements in decoding accuracy, increasing percent correct by 4–6% on average across participants.

### 3.5 GLMdenoise improves brain-representational predictions of perceived object dissimilarity

To further assess whether GLMdenoise provides better estimates of task-related BOLD signals, we exploited behavioral measurements available in one of our experiments. Such measurements are independent of the physiological measurements provided by fMRI, and thus could provide independent validation of the fMRI results. In Experiment 4, participants engaged in a behavioral experiment in which they were asked to arrange the stimuli used in the fMRI experiment according to their similarity using an interactive display (Charest et al., 2014; Kriegeskorte and Mur, 2012). The purpose of this experiment was to characterize each participant’s unique subjective experience of the stimuli and to potentially link these perceptual effects to the participant’s brain representations (Charest et al., 2014). We computed Pearson’s correlation between RDMs computed from a region of interest drawn around the inferior temporal cortex of the participants (brain RDM) and RDMs computed from the behavioral experiment (behavior RDM). This was done for the subset of stimuli that were common across all participants. For nearly every participant (19/20), we observed an increase in correlation between the brain RDM and the behavior RDM when using GLMdenoise to construct the brain RDM (Figure 6). This is an important observation because the denoising performed on the fMRI data has no access to the behavioral similarity structure in any way. The improvement in brain-behavior correlation demonstrates that GLMdenoise improves the accuracy of information extracted from the brain data.

**Figure 6.**
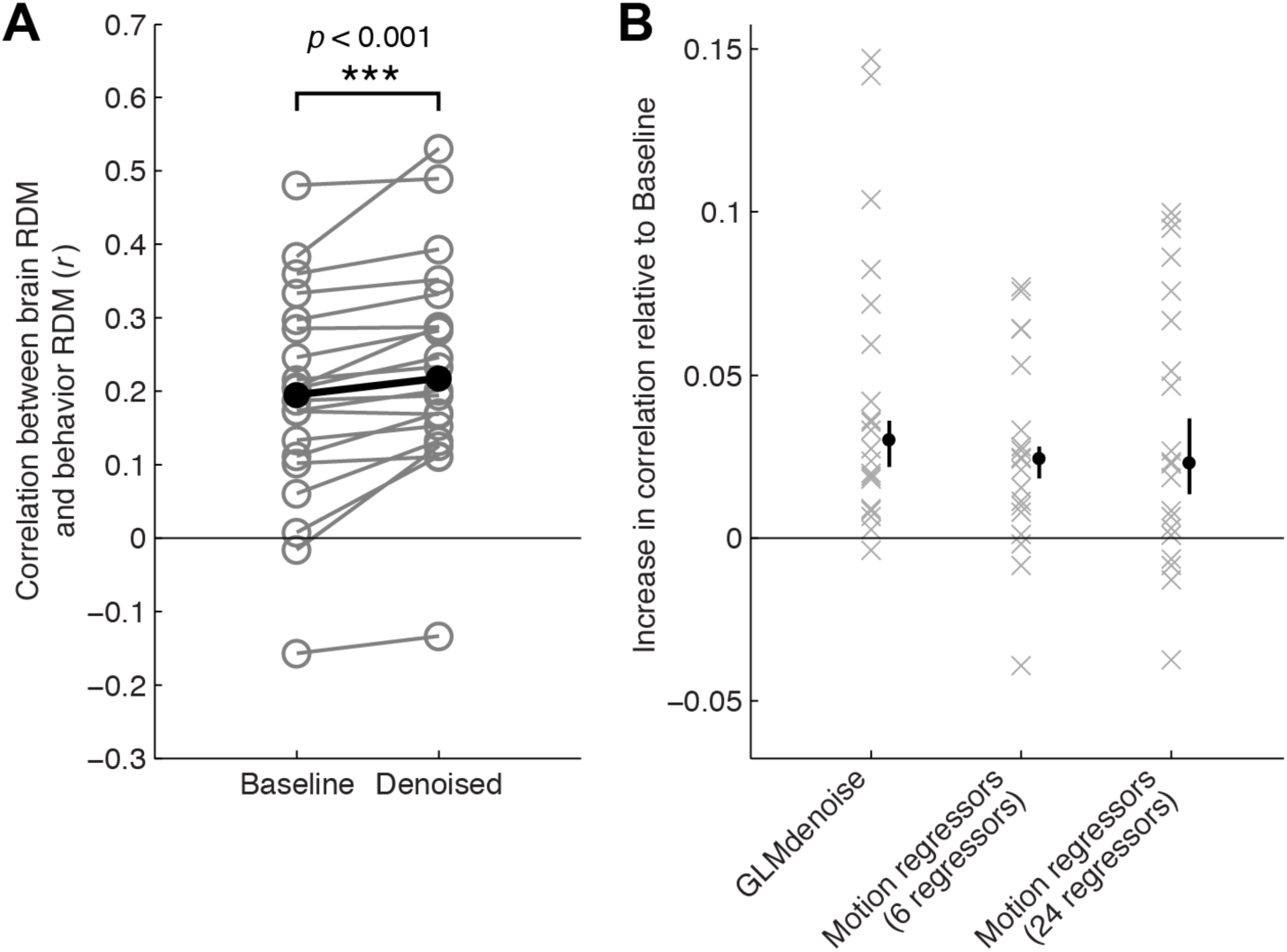
Denoising of brain-based RDMs improves correspondence with behavior-based RDMs. In Experiment 4, participants were asked to make behavioral judgments about the stimuli used in the fMRI experiment, and an RDM was constructed from these behavioral judgments. We averaged the Test and Re-test RDMs derived from the fMRI data and then correlated the lower triangle of this brain-based RDM to the lower triangle of the behavior-based RDM. (A) Results using GLMdenoise. This panel shows the correlation observed for each participant, before and after denoising. Black dots indicate the median across participants. Nearly all (19/20) participants show an increase in correlation and this increase in correlation is statistically significant (two-tailed sign test). (B) Comparison to other approaches. Each ‘x’ indicates for one participant the increase in brain-behavior correlation that is observed compared to Baseline (no denoising). The black dot indicates the median increase across participants (error bars indicate 68% confidence intervals obtained by bootstrapping participants). Although differences in performance across the three approaches are not statistically significant, notice that GLMdenoise consistently improves performance, whereas the Motion regressors approaches yield more mixed results (some datasets improve, some datasets worsen).

## 4. DISCUSSION

### 4.1 GLMdenoise yields more reliable estimates of task-related activity patterns

In this study, we have shown that our earlier observation of improved cross-validation accuracy of beta weights after application of GLMdenoise (Kay et al., 2013a) translates into practical benefits for studies using multivariate analyses. First, representational dissimilarity matrices estimated from independent splits of the data from each participant are more replicable after application of GLMdenoise. This indicates that GLMdenoise improves the reliability of activity pattern estimates and representational geometries. This improved reliability of pattern estimates also translates to greater pattern classification accuracies, stronger correlations between perceptual judgments and brain representations, and improved consistency of representational geometries at the group level.

### 4.2 GLMdenoise estimates and removes a wide variety of sources of nuisance variation

The philosophy behind GLMdenoise is similar to existing strategies for removing noise from neuroimaging data. Several denoising practices exist in which nuisance regressors are considered and removed from the data. Perhaps the most common denoising practice is the inclusion of motion parameters as additional regressors in the general linear model (Bright and Murphy, 2015; Monti, 2011; Pernet, 2014). Other sources of noise that are often included involve auxiliary physiological measurements, such as cardiac and respiratory measurements to predict some of the physiological noise components in the BOLD signal (Birn et al., 2006; Chang et al., 2009; Glover et al., 2000; Hagberg et al., 2012; Shmueli et al., 2007). There are two key advantages offered by GLMdenoise over these existing methods. One is that GLMdenoise captures all of these types of nuisance effects. Another is that GLMdenoise removes nuisance effects in a way that is specifically designed to not overfit the data. For example, as we showed previously (Kay et al., 2013a) and as shown in the present study (see Figure 4A), including motion parameters in the linear model often does help, but also has the capacity to hurt. As a matter of design, GLMdenoise optimizes the number of noise regressors used on each dataset. This is important because whether modeling out nuisance effects is effective depends on the magnitude of the nuisance effects, the magnitude of the task-related signals, and the amount of correlation between the nuisance effects and task-related signals, all of which may depend on the participant and the experiment (Kay et al., 2013a). We clarify the nature of these contingencies using a set of simple simulations (Supplementary Figure 2). These simulations demonstrate that modeling task-correlated nuisance effects improves model accuracy when nuisance effects are strong and highly correlated with the task, but can degrade model accuracy when nuisance effects are weak. The latter scenario might occur if one blindly includes nuisance regressors into an fMRI design matrix (due to overfitting); GLMdenoise guards against this possibility by assessing model accuracy through cross-validation.

One possible concern with using GLMdenoise is that it somehow leads to fMRI beta weights that do not reflect the ‘true’ activity patterns elicited by the experimental conditions. For example, if a denoising method altered beta weights by biasing them towards the mean beta weight, this would improve replicability at the expense of pulling condition-specific activity patterns away from the true underlying activity patterns. Such bias could potentially underlie the observation that denoised RDMs are smoother than baseline RDMs (see Figure 1). There are three considerations that argue against this possibility. First, considering the nature of the technique, we see that there is no explicit mixing of signals across voxels (aside from the fact that the nuisance regressors are derived from a common noise pool): each voxel is independently modeled by the GLM and there is no restriction on the weights associated with the nuisance regressors for each voxel. Thus, it is difficult to see how some sort of smoothing bias could result from the GLMdenoise procedure. Second, in our RDM analysis, we found that undenoised RDMs are better predicted, using an independent split of the data, by denoised RDMs than by undenoised RDMs. This suggests that denoising does not induce bias but instead reduces variance. Third, we exploit the similarity judgments collected in Experiment 4 as an external validation of brain RDM estimates. We were able to confirm, using this different measurement modality, that denoising brain measurements improves the correspondence between perceived similarity and the brain’s similarity structure. Given these considerations, we suggest that GLMdenoise provides substantially better estimates of task-related activity patterns without inducing appreciable bias.

### 4.3 What preconditions and complications are associated with the application of GLMdenoise?

The GLMdenoise technique is general (it requires only a design matrix and fMRI time-series) and fully automated (it requires no hand-tuning of parameters, although it can be customized if desired). Furthermore, because no physiological recordings nor additional fMRI data are required, the technique can be retrospectively applied to existing fMRI datasets. Despite these appealing features, it is important to recognize some caveats and limitations:

- Since GLMdenoise relies on cross-validation of task-related BOLD signals, GLMdenoise is not applicable to resting-state fMRI.
- GLMdenoise requires multiple fMRI runs, with presentation of each condition more than once. This is necessary because GLMdenoise involves cross-validation across runs. Conventional denoising techniques are recommended for experiments with only one repetition per condition.
- Because GLMdenoise is fully data-driven, the nature of the noise removed is unclear without further analyses. Moreover, the amount of noise removed and its properties may vary across experiments and participants.
- A central assumption of GLMdenoise is that the fMRI measurements (including both task-related signals and noise sources) are relatively stationary across runs. In other words, evoked BOLD signals should be replicable across runs and general trial distributions should be relatively balanced throughout the experiment.
- The number of noise regressors in GLMdenoise is, by default, optimized with respect to all voxels exhibiting task-related signals. It is possible that specific brain regions might degrade in accuracy while the rest of the dataset improves. If one desires, one can restrict the optimization to specific voxels of interest. This and other customizations are implemented in the GLMdenoise toolbox.

Bearing in mind these caveats and limitations, we believe that the substantial improvements in the quality of MVPA results we have demonstrated make GLMdenoise a valuable analysis tool.

## 5. AUTHOR CONTRIBUTIONS

I.C. and K.K. conducted the experiments, analyzed the data, and wrote the paper. All authors discussed the results and edited the manuscript.

## 6. ACKNOWLEDGMENTS

We thank J. Winawer for helpful comments. This work was supported by an European Research Council (ERC) Starting Grant ERC-2017-StG 759432 (to I.C.), by an UK Medical Research Council Grant MC-A060-5PR60 and an ERC Starting Grant ERC-2010-StG 261352 (to N.K.), and by the McDonnell Center for Systems Neuroscience and Arts & Sciences at Washington University (to K.K.).

## 7. COMPETING INTERESTS

The authors confirm that there are no competing interests.

## 8. APPENDIX A. SUPPLEMENTARY DATA

Supplementary data associated with this article can be found, in the online version, at doi:XXX.

---

**Supplementary Figure 1.**
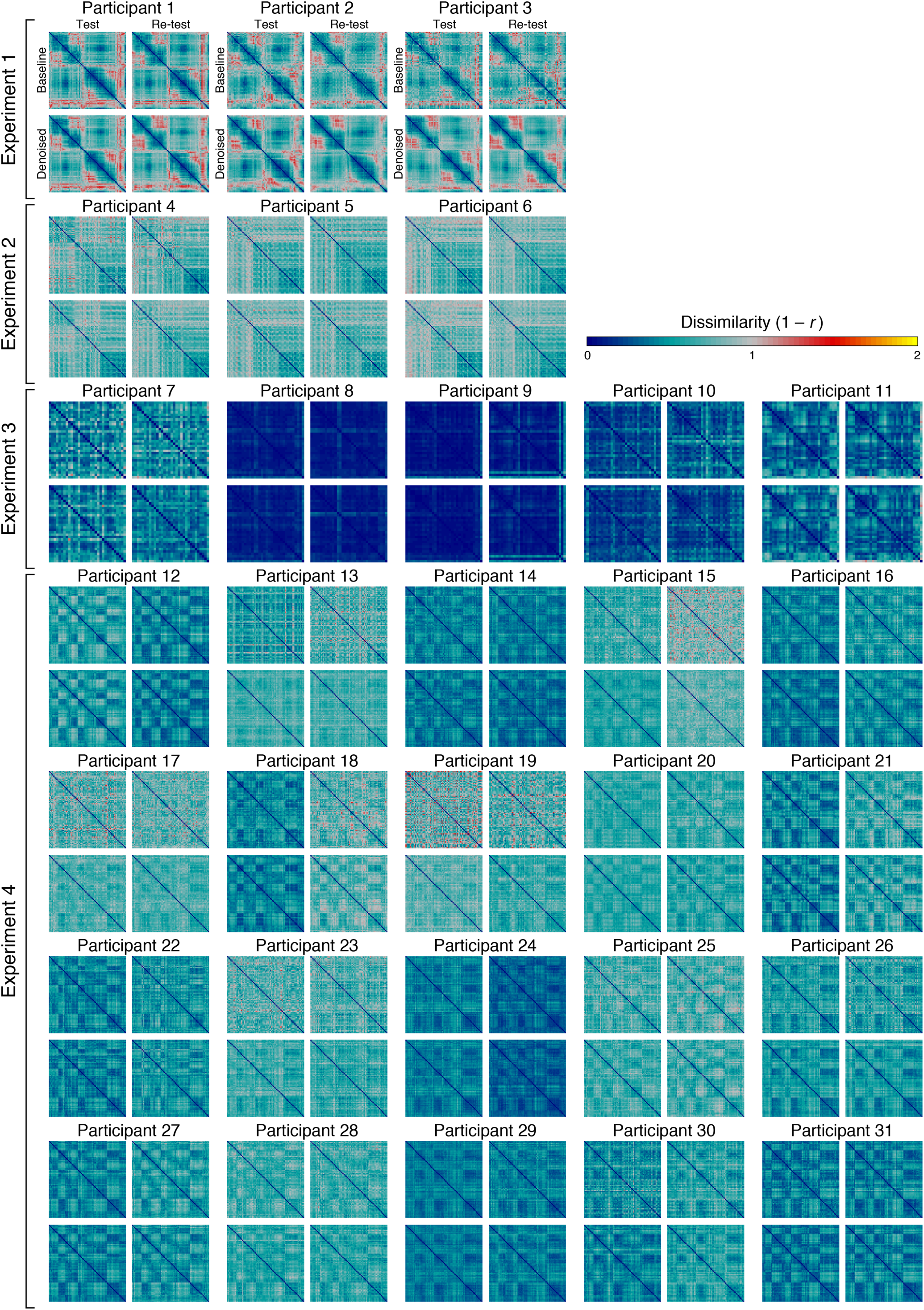
RDM results for all participants. Same format as Figure 1.

**Supplementary Figure 2.**
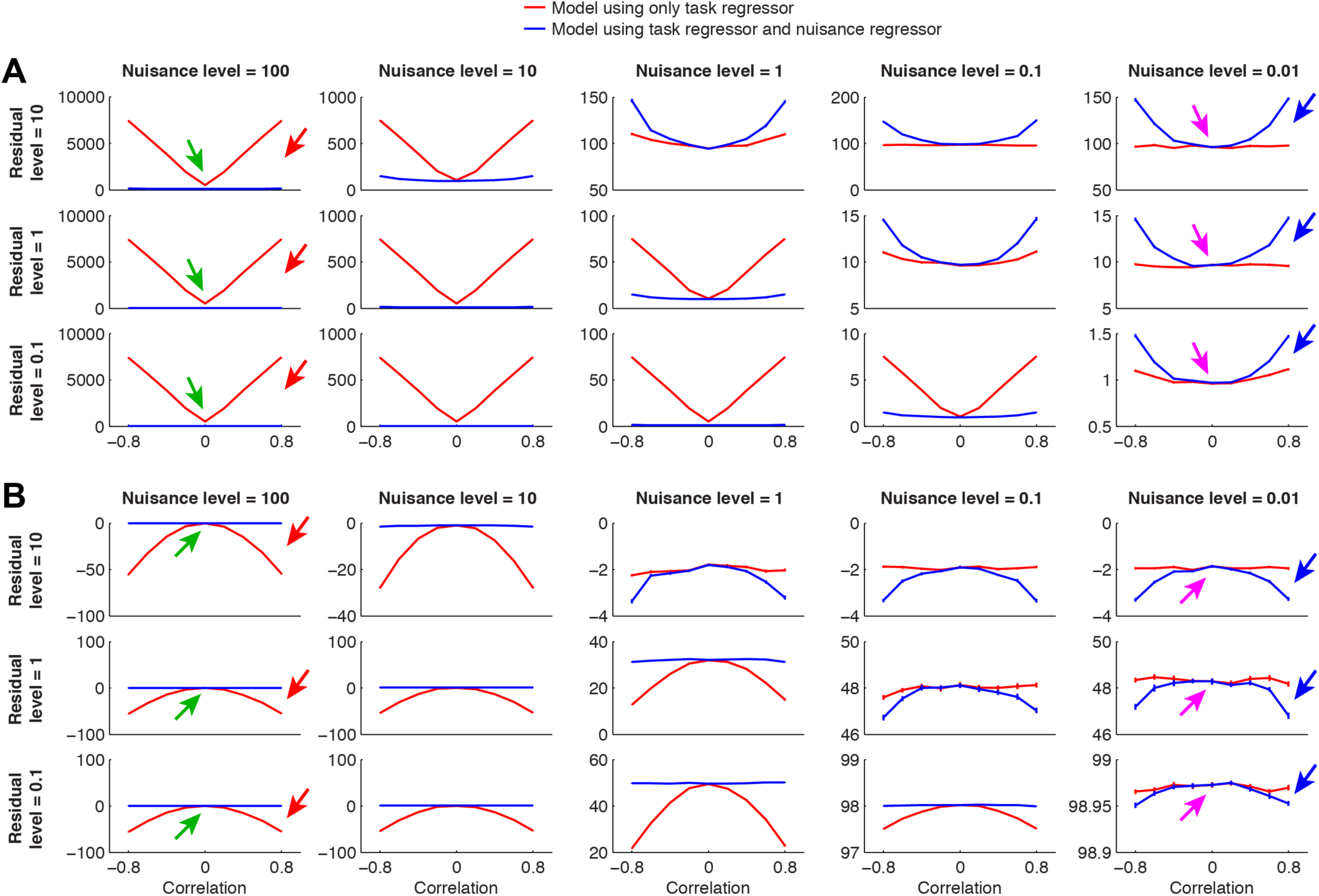
Modeling task-correlated nuisance effects can improve model accuracy. An assumption of GLMdenoise is that task-related fluctuations and nuisance fluctuations are uncorrelated in the sense that the expected value of the correlation between the two types of fluctuations is zero. However, in any given limited sample of data, the correlation between task-related and nuisance fluctuations may be non-zero, and this phenomenon is what GLMdenoise exploits to improve the accuracy of task estimates. To illustrate, we perform simulations (see Methods for details; simulation code available at http://osf.io/bf736) in which we randomly generate task and nuisance regressors, simulate time-series data that reflect a mixture of task, nuisance effects, and residuals, and then fit models to the time-series data in an attempt to estimate the task-related component of the data. We evaluate two models: one model consists of only the task regressor (red line) and the other model consists of both the task regressor and the nuisance regressor (blue line). Here we show simulation results in which we systematically vary the nuisance level, as depicted by the columns, and the residual level, as depicted by the rows. In each plot, the *y*-axis indicates the absolute value of the difference between the estimated and true task weight (panel A) or the cross-validated *R*^2^ of the task estimate (panel B). Results are binned according to the correlation observed between the task regressor and the nuisance regressor, as depicted by the *x*-axis. For each bin, the median across 10,000 simulations and the 68% confidence interval on the median (bootstrap procedure) is shown. The results indicate that when the nuisance effects are strong, inclusion of the nuisance regressor improves performance when the task and nuisance regressors are correlated (red arrows) but does not affect performance when the task and nuisance regressors are uncorrelated (green arrows). In contrast, when the nuisance effects are weak, inclusion of the nuisance regressor degrades performance when the task and nuisance regressors are correlated (blue arrows) but does not affect performance when the task and nuisance regressors are uncorrelated (magenta arrows). Thus, these results demonstrate that modeling task-correlated nuisance effects *can* improve task estimates, but this is contingent on the strength of the nuisance effects and the level of correlation that exists between the task and nuisance regressors. GLMdenoise uses cross-validation to determine on a case-by-case basis whether including nuisance regressors improves or degrades task estimates.

## Footnotes

1 Note that ‘noise’ is operationally defined as signal fluctuations that are not captured by the GLM design matrix, and so the noise regressors may include genuine, neurally-driven BOLD signals that are not of interest to the experimenter.

